## Supplemental Materials for "Homeostatic Control of Deep Sleep in *Drosophila*: Implications for Discovering Correlates of Sleep Pressure"

*Running title: Drosophila sleep homeostasis*

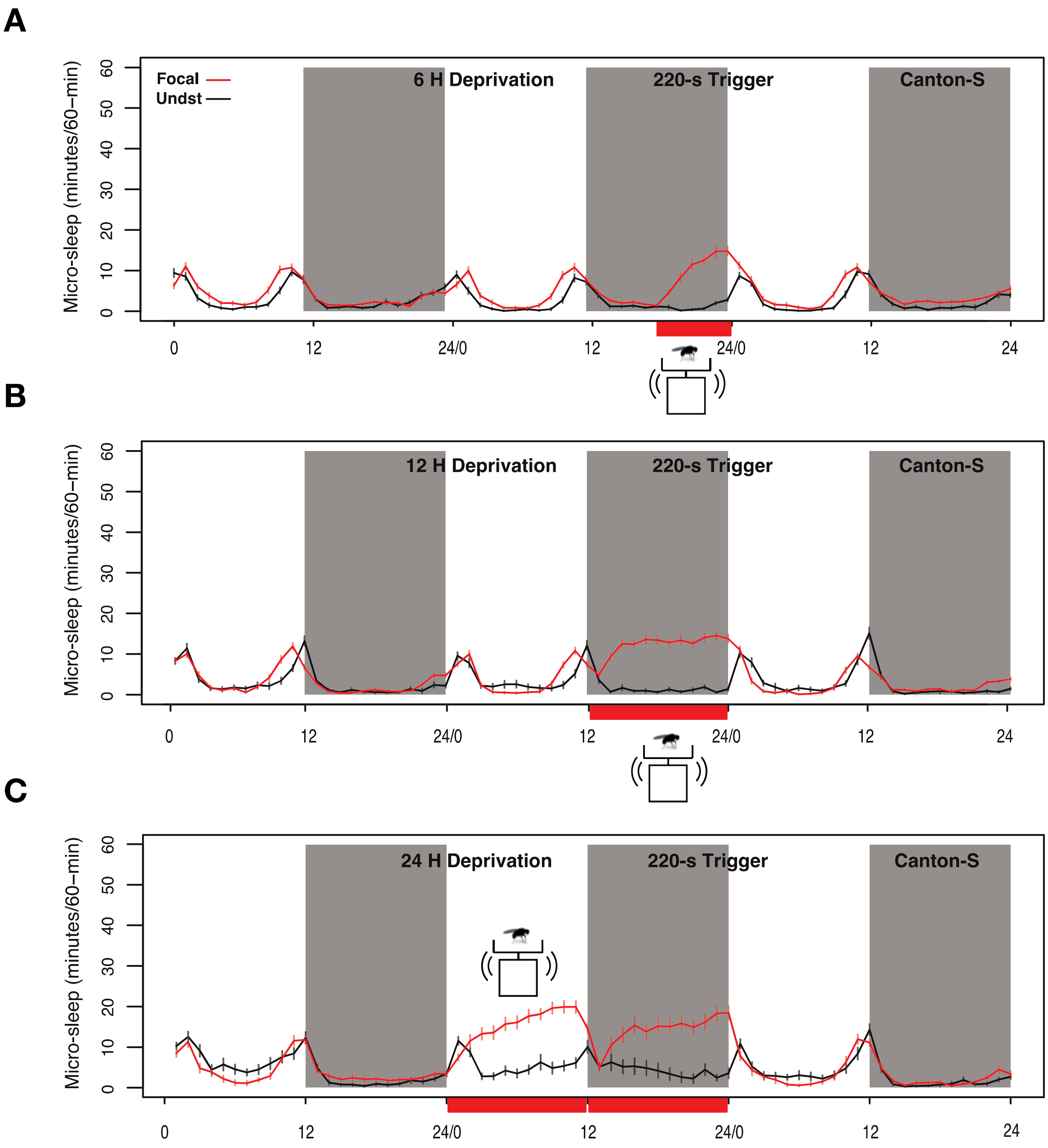


**Supplementary Figure 1:** Time courses of “active sleep” – bouts of inactivity that are between one to five minutes long – under conditions of 6 **(A)**, 12 **(B)** and 24-h **(C)** sleep deprivation using vortexers at a trigger frequency of 220-s for *CS* flies. Plots are means±SEM. Gray shaded regions indicate the dark phase of the LD cycle. Red shaded regions along the *x*-axis indicate windows of sleep deprivation.

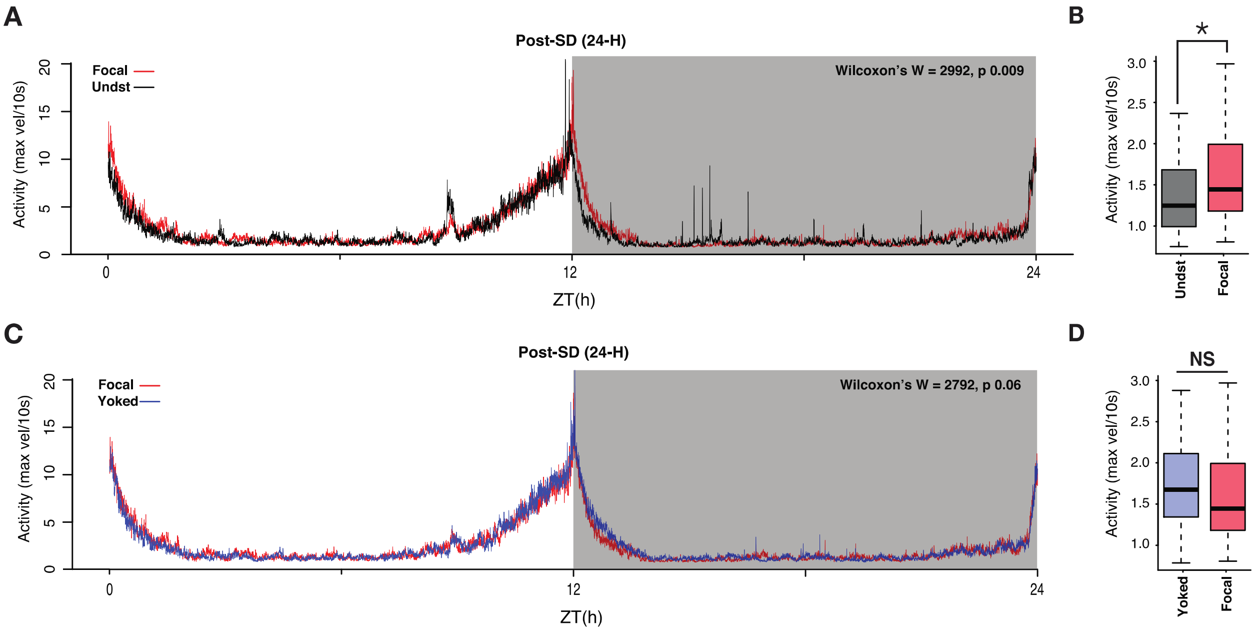


**Supplementary Figure 2. A.** Activity time course of undisturbed and sleep-deprived flies across the 24 hours following mechanical deprivation. During ZT12-24 on post SD day 1, focal flies display elevated night-time activity at times corresponding to the normal period of consolidated sleep. **B**. The activity of focal flies between ZT12-24 was significantly higher than that of undisturbed controls (Wilcoxon's $W = 2902$, $p = 0.009$). **C.** Activity time course of focal and yoked flies during the same Post deprivation window (24-H). **D.** Activity of focal and yoked flies during ZT12-24 were not significantly different from each other (Wilcoxon's $W = 2902$, $p = 0.06$). $n = 82$ each for focal and yoked categories, and $n = 58$ for unperturbed controls.

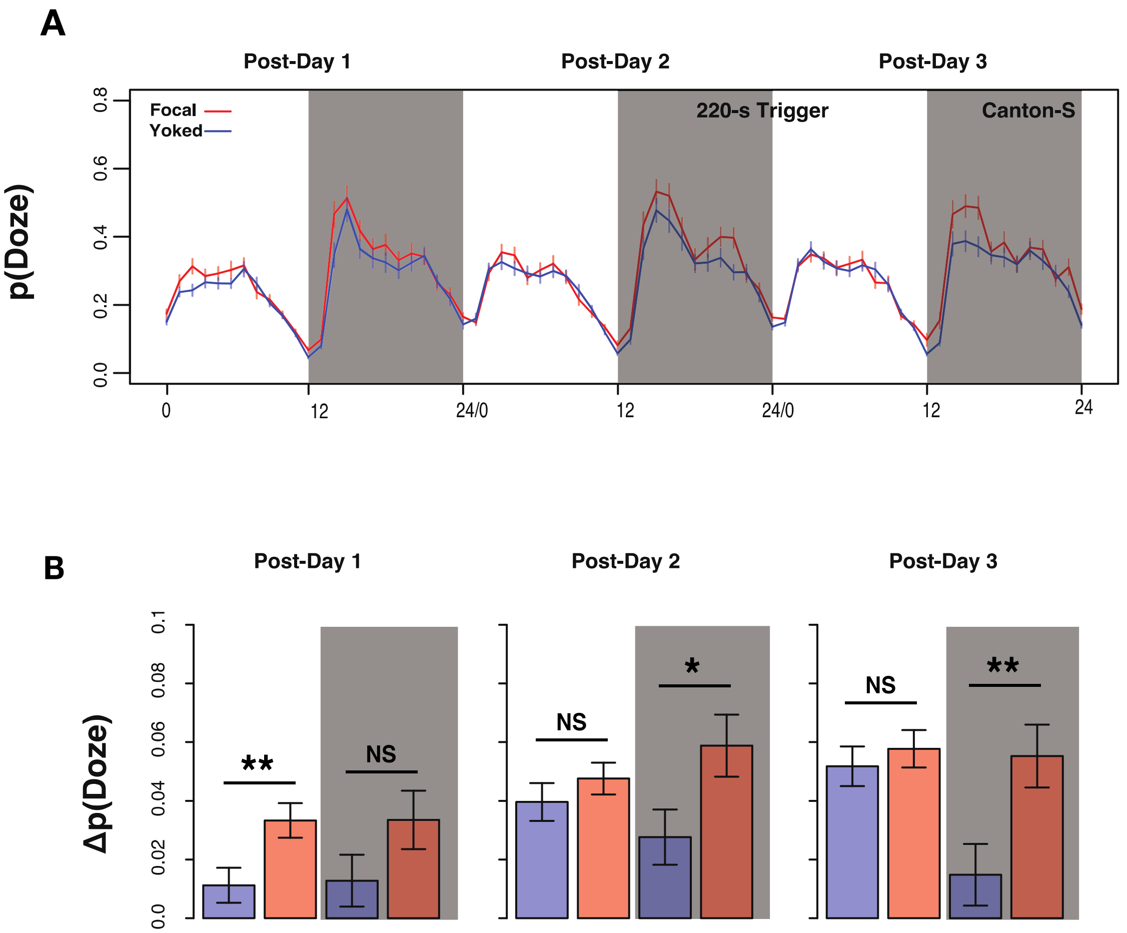


**Supplementary Figure 3. A.** Time course of conditional Doze probability – p(Doze) – for focal and yoked *CS* controlled flies across three cycles post sleep deprivation using 220-s trigger frequency. Plotted are means±SEM. **B.** Changes in p(Doze) values compared to baseline. Plotted are means±SEM. Statistically significant differences between focal and yoked flies were inferred using a two-sample, one tailed t-tests for unequal variances (Post1: Day – $t_{161.99} = 2.64$, $p = 0.005$; Night – $t_{159.8} = 1.56$, $p = 0.06$; Post2: Day – $t_{157.66} = 0.95$, $p = 0.17$; Night – $t_{159.83} = 2.2$, $p = 0.01$; Post3: Day – $t_{161.41} = 0.64$, $p = 0.26$; Night – $t_{161.93} = 2.7$, $p = 0.004$). Gray shaded regions indicate dark phase of the LD cycle, in both panels. * $p < 0.05$, ** $p < 0.01$, NS – Not Significant.

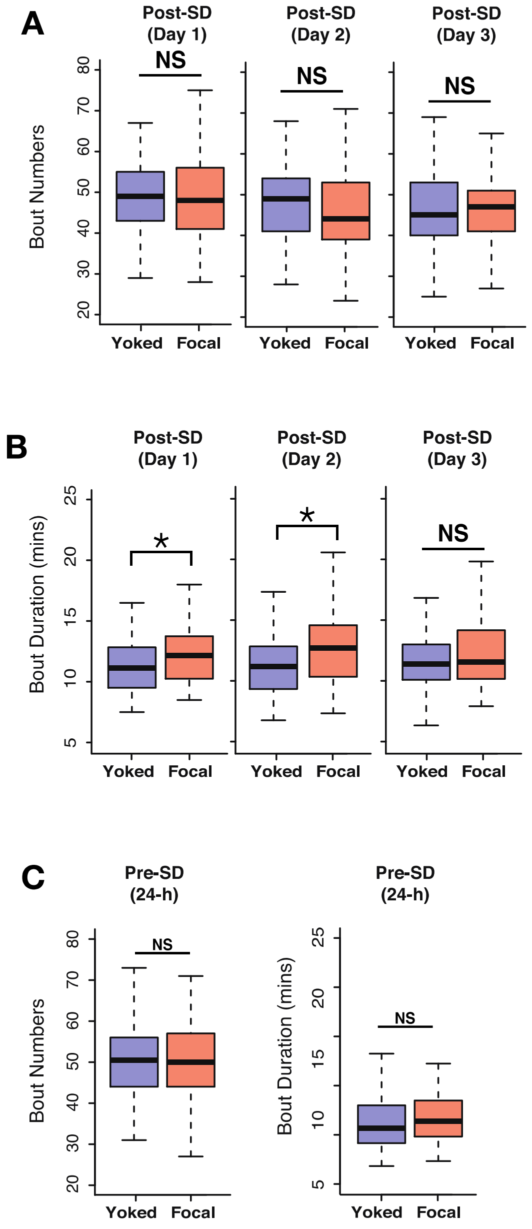


**Supplementary Figure 4. A.** Comparison of bout numbers for standard sleep between focal and yoked flies in three post-deprivation cycles did not reveal significant differences (Post1: Wilcoxon's $W = 3385$, $p = 0.94$; Post 2: Wilcoxon's $W = 2962.5$, $p = 0.18$; Post 3: Wilcoxon's $W = 3273$, $p = 0.77$). **B.** Comparing bout duration of standard sleep across three post deprivation cycles revealed that focal flies had significantly higher bout durations for the first two recovery cycles (Post1: Wilcoxon's $W = 4077$, $p = 0.01$; Post 2: Wilcoxon's $W = 4269$, $p = 0.002$; Post 3: Wilcoxon's $W = 3624$, $p = 0.38$). $n = 82$ each for focal and yoked categories. Undisturbed control n = 58 * < 0.05, and N.S. (Not Significant). **C.** Pre-deprivation bout numbers (Wilcoxon's $W = 3358$, $p = 0.99$) and bout durations (Wilcoxon's $W = 3705$, $p = 0.25$) were not significantly different between focal and yoked pairs). $n = 82$ each for focal and yoked categories, and n = 58 for unperturbed controls * < 0.05, and N.S. (Not Significant).

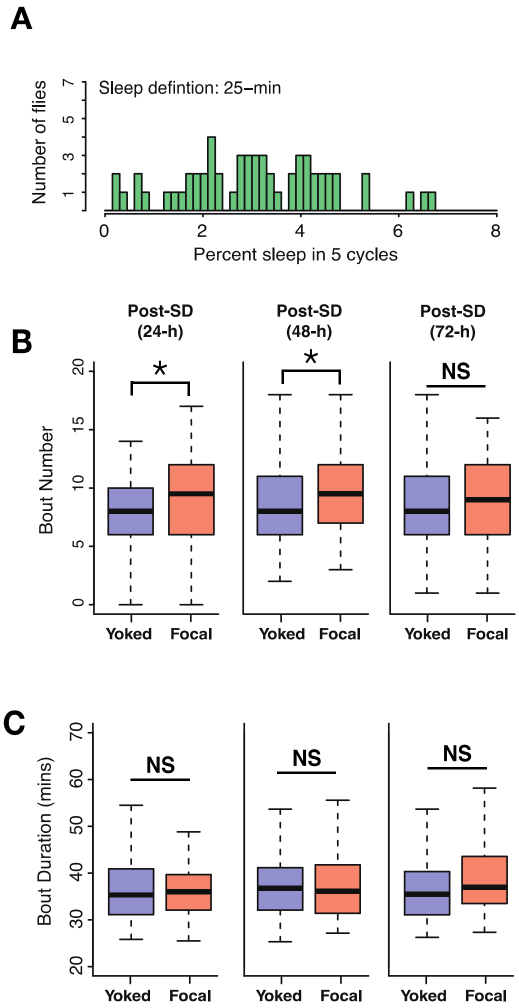


**Supplementary Figure 5. A.** frequency distribution of 25-min sleep bouts in undisturbed flies collected from 5 cycles. Every fly displayed at least one bout of sleep that was 25-min or longer. **B.** Total bout numbers in 25-min, long bout sleep across two cycles following 24-h of deprivation using 220-s inactivity triggers showed significant increases in focal flies compared to yoked flies (Post1: Wilcoxon's $W$ = 4029, $p$ = 0.02; Post 2: Wilcoxon's $W$ = 4186.5, $p$ = 0.006; Post 3: Wilcoxon's $W$ = 3779.5, $p$ = 0.16). **C.** Duration of long bouts are not significantly different between focal and yoked flies in three post-deprivation days (Post1: Wilcoxon's $W$ = 3422, $p$ = 0.53; Post 2: Wilcoxon's $W$ = 3229.5, $p$ = 0.66; Post 3: Wilcoxon's $W$ = 3888, $p$ = 0.08). $n$ = 82 each for focal and yoked categories, and n = 58 for unperturbed controls * < 0.05, and N.S. (Not Significant).

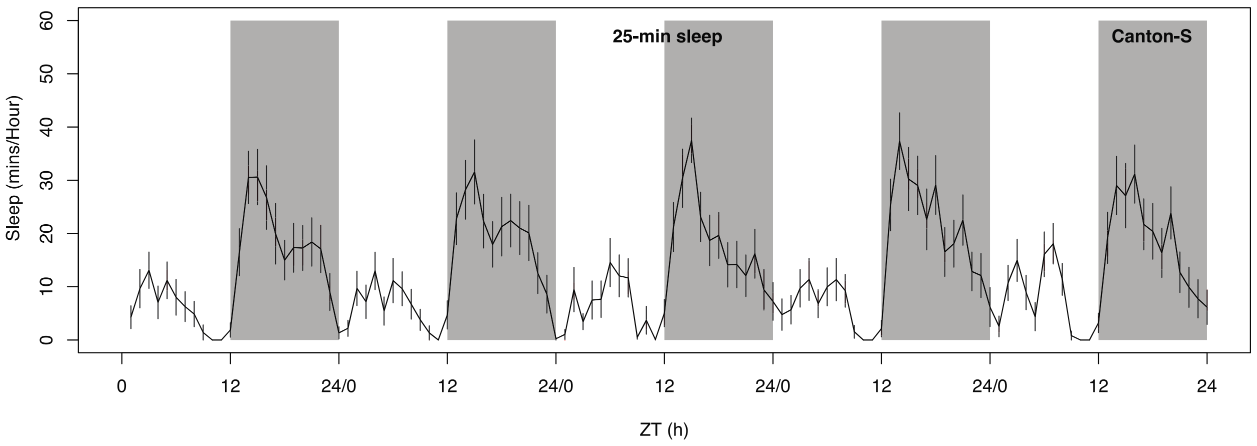


**Supplementary Figure 6.** Time course of control *CS* flies for five LD cycles from a different run (see and compare with Figure 4), showing the remarkable consistency between runs in the amount of 25-minute sleep. Plotted are means±SEM and dark shaded regions indicate the dark phase of the LD cycle.

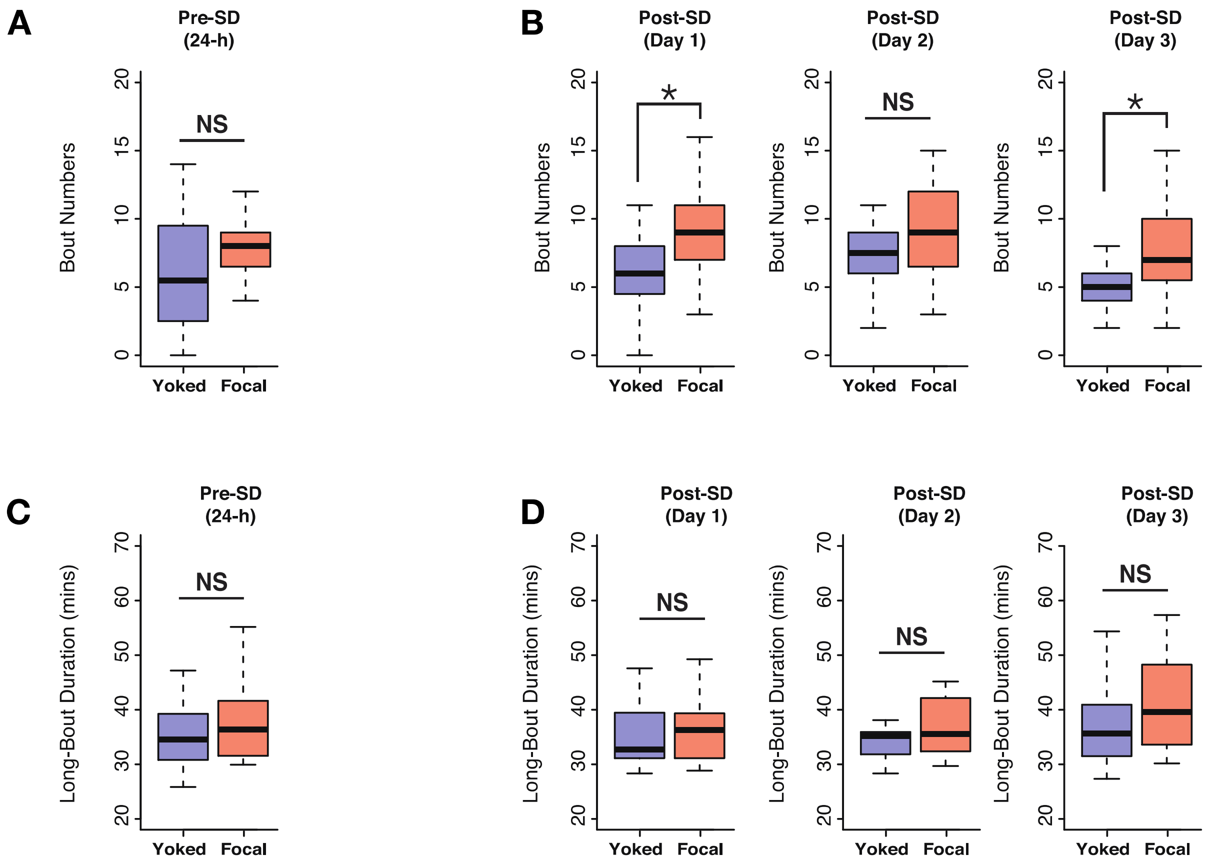


**Supplementary Figure 7. A.** Long-sleep bouts prior to deprivation were not different between focal and yoked flies (Wilcoxon's $W$ = 263, $p$ = 0.09). **B.** On post-deprivation day 1 focal flies showed significantly increased long bout numbers (Wilcoxon's $W$ = 304, $p$ = 0.004). On second day even though a trend was seen, it was not significant (Wilcoxon's $W$ = 244.5, $p$ = 0.23), before returning to significant levels on post-deprivation day 3 (Wilcoxon's $W$ = 295, $p$ = 0.009). **C.** Long bout duration on baseline days was not different between focal and yoked flies (Wilcoxon's $W$ = 225.5, $p$ = 0.32). **D.** On three post derivation days, focal and yoked pairs did not show significant differences in long bout durations (Post1: Wilcoxon's $W$ = 210.5, $p$ = 0.57; Post 2: Wilcoxon's $W$ = 231.5, $p$ = 0.24; Post 3: Wilcoxon's $W$ = 242.5, $p$ = 0.14). $n$ = 20 each for focal and yoked categories. * < 0.05, and N.S. (Not Significant).

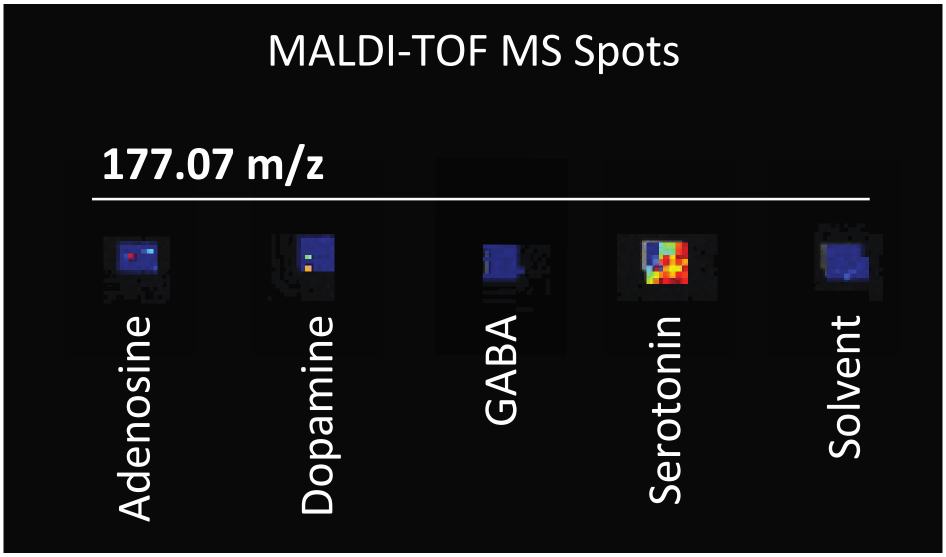


**Supplementary figure 8.** Spotted amines in MALDI-TOF MS experiments and detected serotonin concentration at m/z 177.07. Water was used as a solvent to make 1mg/ml concentrations for spotting slides.
